# Inhibition of EV71 replication by an interferon-stimulated gene product L3HYPDH

**DOI:** 10.1101/304345

**Authors:** Jian Liu, Luogen Liu, Shinuan Zeng, Xiaobin Meng, Nanfeng Lei, Hai Yang, Runcai Li, Xin Mu, Xuemin Guo

## Abstract

Enterovirus 71 (EV71) is the common causative agent of hand-foot-mouth disease (HFMD). Despite evidence in mice model suggested that the interferon (IFN) signaling pathways play a role in defending against this virus, knowledge on the IFN-mediated antiviral response is still limited. Here we identified an IFN-stimulated gene (ISG) called L3HYPHD, whose expression inhibits EV71 replication. Mapping assay indicated that amino acids 61-120 and 295-354 are critical for its optimal antiviral activity. Mechanismly, L3HYPDH specifically inhibits protein translation mediated by EV71 internal ribosome entry site (IRES). Our data thus uncovered a new mechanism utilized by the host cell to restrict EV71 replication.

## 1. Introduction

Hand-foot-mouth disease (HFMD) affects infants and children across the Asian-Pacific region, characterized by fever, rash, and occasionally severe neurological symptoms [1]. Its major causative agent is enterovirus 71 (EV71), an enterovirus belonging to the Picornaviridae family. RNA is used as the genomic material, single and positive-stranded, with a length of about 7,400 nt. The genomic RNA encodes only one open reading frame (ORF) flanked with a 5’-untranslated region (5’-UTR) and a 3’-UTR [2,3]. The 5’-UTR contains a cloverleaf structure and an internal ribosome entry site (IRES), responsible for viral RNA replication and protein translation, respectively [4]. The life cycle of EV71 starts with attachment to the host cell surface by recognition of specific receptors, followed by endocytosis and release of viral RNA into the cytoplasm [5]. Then, EV71 IRES initiates viral protein translation by recruiting translation machineries. The polyproteins are processed into structural and non-structural proteins by the protease 2A and 3C embedded. When viral proteins accumulate, viral protein 3CD binds to the cloverleaf structure of 5’-UTR to initiate viral RNA replication. The newly synthesized plus-stranded RNAs in turn direct viral protein synthesis in large quantities. With the assembly of viral RNAs and proteins into virions, the host cell lyses and progeny viruses are released for a new round of infection [6].

EV71 is capable to suppress type I interferon (IFN-I) production and its signaling activation [7–10]. Nonetheless, cells infected with EV71 can still respond to type I IFN treatment and display an enhanced antiviral state. For example, in vitro studies showed that some type I IFNs, including IFN-α4, IFN-α6, IFN-α14 and IFN-α16, significantly reduced cytopathic effect (CPE) induced by EV71 infection [11]. An IFN-α2b aerosol therapy has been used topically to treat HFMD [12]. We recently reported an IFN-induced gene TMEM106A encodes proteins to inhibit EV71 infection through restricting viral binding onto host cells [13]. However, how IFNs suppress EV71 infection is still largely unknown.

Many new interferon-stimulated genes (ISGs) were identified from human immune cell lines after treatment with interferon [14], but many of them are unclear of the antiviral activities. Using a fluorescent activated cell sorting-based strategy for screening, we identified several ISGs with anti-EV71 efficacy (data not shown). One of them is C14orf149, which was identified as a gene encoding a trans-3-hydroxy-L-proline dehydratase and then renamed L3HYPDH [15]. We found that L3HYPDH, possesses antiviral activity against EV71, and its mechanism of action was investigated with a series of biochemical and genetic assays.

## 2. Materials and Methods

### Plasmids construction

pCAG-DsRed, a red fluorescent protein-expressing plasmid, has been described previously [16]. pWSK-EV71-GFP is an infectious EV71-GFP cDNA clone, with a GFP-coding sequence inserted downstream of EV71 5’UTR and in frame fusion with the downstream VP4, and expression of EV71-GFP is driven by a T7 promoter [17]. pcDNA3.1-T7RNP expresses T7 RNA polymerase. These plasmids were kindly provided by Dr. Liguo Zhang at the Institute of Biophysics, Chinese Academy of Sciences (IBP, CAS). A siRNA targeting the coding sequence of L3HYPDH from position 791 to 811 was designed according to the recommendation of Sigma-Aldrich (https://www.sigmaaldrich.com/catalog/genes) and named shRNA149. A pair of complementary oligonucleotides 5’-GATCCCCCAGATGAACAGGTTGACAGAATTCAAGAGATTCTGTCAACCTGTTCATCTGTTTTTA-3’ (sense) and 5’-AGCTTAAAAACAGATGAA CAGGTTGACAGAATCTCTTGAATTCTGTCAACCTGTTCATCTGGGG-3’ (antisense) were synthesized with 5’ ends being *BglII* and *HindIII* restriction site overhangs. For each oligonucleotide, the target sequence was sense followed by antisense orientations separated by a nine-nucleotide spacer. Oligonucleotides were annealed and then cloned into the *BglII* and *HindIII* sites of pSUPER.retro.neo+gfp (Oligoengine, herein abbreviated for pSUPER-GFP) to generate pSUPER-GFP-shRNA149. L3HYPDH wild type (WT) and deletion mutants as indicated in Figure 3 were amplified with PCR using pLPCX-C14orf149 (L3HYPDH) [14], kindly provided by Dr. Guangxia Gao at IBP, CAS, as the template. PCR products of L3HYPDH WT and deleted mutants were digested with *BamHI* & *NotI* and *KpnI* & *XbaI*, respectively, and inserted into similarly digested pcDNA4-To/myc-His B (Invitrogen), resulting in pcDNA4-L3HYPDH, pcDNA4-L3HYPDHΔN1, pcDNA4-L3HYPDHΔN2, pcDNA4-L3HYPDHΔN3, pcDNA4-L3HYPDHΔC1, pcDNA4-L3HYPDHΔC2, and pcDNA4-L3HYPDHΔC3. psiCHECK2-M was a modified form of psiCHECK-2 (Promega) with deletion of the HSV-TK promoter (Figureure 5A). Inverse PCR was performed with high-fidelity DNA polymerase Phusion (ThermoFisher) and a pair of back-to-back primers to amplify the whole plasmid except the HSV-TK promoter sequence. PCR products were self-ligated and resulted in psiCHECK2-M; meanwhile, a *SalI* and a *NotI* sites within the back-to-back primers were introduced into the plasmid. EV71-5’UTR and HCV (Hepatitis C virus)-5’UTR were amplified from pWSK-EV71-GFP and pNL4-3RL-HCV-FL [18] by PCR, respectively. After digestion with *SalI* and *NotI*, the PCR products were linked into the similarly digested psiCHECK2-M and resulted in psiCHECK2-M-EV71-5’UTR and psiCHECK2-M-HCV-5’UTR. All primers used are listed in Table S1.

### Cell culture and virus preparation

293A, 293A-SCARB2, RD, Vero, HeLa, and A549 cells were cultured in DMEM (Gibco) supplemented with 10% fetal bovine serum (FBS, Gibco). 293A-SCARB2 (Kindly provided by Dr. Liguo Zhang at IBP, CAS), originated from a 293A cell line and constitutively expresses the main EV71 receptor scavenger receptor class B member 2 (SCARB2). To generate the cell line constitutively expressing tagged L3HYPDH, 293A-SCARB2 cells were transfected with pcDNA4-L3HYPDH as described below and selected with Zeocin (200 μg/ml). Resistant colonies were individually expanded and detected by Western blot. One positive clone was chosen and named 293A-SCARB2-L3HYPDH. This process was applied to the empty vector and resulted in control cell 293A-SCARB2-Ctrl.

EV71-MZ (GenBank accession no. KY582572), isolated from the throat swab of an ICU patient at Meizhou People’s Hospital in 2014 [19], was amplified by successive passages in RD cells until apparent CPE appeared. EV71-GFP was generated by co-transfecting pWSK-EV71-GFP and pcDNA3.1-T7RNAP into 293A-SCARB2 cells as described previously [17]. Viral supernatants were titrated using a plague assay, aliquoted, and then used for infection.

### Transfection and infection

Depending on the experiments, cells were seeded into a 24-well or 6-well plate or 10 cm dish and were grown to approximately 80% confluence prior to transfection or infection. All plasmid and RNA transfections were carried out by using Lipofectamine TM 2000 (Life Technology) according to the manufacturer’s instructions. After incubation for the indicated time, cells were treated as required.

Viral infection was performed by incubating cells with EV71-GFP or EV71-MZ at a different multiplicity of infection (MOI) for 1 h, with shaking every 15 min, and then the unbound viruses were aspirated. Cells were washed with PBS, added fresh medium, and incubated for specific time, followed by FACS assay, RT-qPCR measurement, or supernatant titration.

### Plaque assay

The plaque assay was performed as described previously [20]. Briefly, RD cells were incubated with viral supernatants undiluted or diluted in 10-fold series for 1 h. Subsequently, the supernatants were aspirated, and cells were covered with DMEM containing 1% methylcellulose (Sigma-Aldrich) and 2% FBS. After incubation for 4 days, cells were fixed with 4% paraformaldehyde (Sigma-Aldrich) and stained with 0.1% crystal violet. Plaques were then quantified via visual scoring.

### Fluorescent activated cell sorting (FACS) assay

To measure GFP production from EV71-GFP, 1×106 infected cells were collected and fixed in 4% paraformaldehyde for 15 min. After washing three times with PBS, cells were resuspended in 0.5 ml of PBS for flow cytometry (LSRFortessa, BD). To assess effects of L3HYPDH knockdown by RNAi, the cells were transfected with pSUPER-GFP-shRNA149 or pSUPER-GFP. After incubation for the indicated time, the cells were harvested and washed with PBS. GFP-positive cells were obtained through FACS, and then lysed for Western blot or seeded into a 24-well plate for EV71-GFP infection or reporter plasmid transfection as required.

### IFN stimulation

Cells were treated with 1000 IU/ml of recombinant human IFN-α2b (Prospec) for different time, and then total RNAs were isolated and used to measure specific mRNA abundance by RT-qPCR.

### *In vitro* transcription of EV71-GFP and microscope assay of GFP

pWSK-EV71-GFP was linearized XbaI and EV71-GFP RNAs were transcribed using the T7 RiboMax kit (Promega). After transfection into 293A-SCARB2-L3HYPDH and 293A-SCARB2-Ctrl cells, the GFP signal was observed under a fluorescence microscope (System Microscope BX63, Olympus) at the indicated times; total RNAs were isolated for RT-qPCR assay.

### RNA isolation and RT-qPCR

Total RNAs were isolated from cells using TRI Reagent (Sigma-Aldrich) according to the manufacturer’s instructions. RT-qPCR was carried out as described previously [21]. Briefly, RNAs were treated with DNase using an RQ1 RNase-Free DNase Kit (Promega); cDNAs were synthesized using PrimeScript RT reagent Kit (Takara, Dalian) and then diluted and subjected to quantitative PCR using TransStart Green qPCR SuperMix (TransGen Biotch) in a CFX96 Touch Real-Time PCR Detection System (Bio-Rad). Primers used appear in Table S1.

### Immunofluorescence assay (IFA)

Subcellular localization of L3HYPDH proteins and the attachment and endocytosis of EV71 virions were detected using IFA as described previously [22] with some modifications. Briefly, 293A-SCARB2-L3HYPDH and 293A-SCARB2-Crtl were individually seeded onto a coverslip. Polyclonal antibody (pAb) specific to c-myc (Sigma-Aldrich, 1:100) and Alexa Fluor 555-labeled anti-rabbit IgG (ThermoFisher, 1:100) were used as primary and secondary antibody, respectively, to localize the subcellular distribution of the tagged L3HYPDH proteins. Similarly, 293A-SCARB2-L3HYPDH and 293A-SCARB2-Crtl cells were infected with EV71-MZ (MOI, 100) at 4°C for 1 h to allow viral attachment or incubated for an additional 30 min at 37°C to allow viral endocytosis. The Anti-EV71 VP2 monoclonal antibody (mAb) (Millipore, 1:50) and Alex flour 555-labeled anti-mouse IgG (ThermoFisher, 1:100) were used as primary and secondary antibody, respectively, to visualize EV71 virions. Nuclei were stained with DAPI (Roche). Fluorescent images of cells were captured using a Zeiss LSM780 META confocal imaging system.

### Western blot

Western blot was performed as described previously [22] with some modifications. Briefly, 48 h after transfection, the cells were lysed with SDS-lysis buffer (30 mM SDS, 50 mM pH 6.8 Tris-HCL, 100 mM DTT and 20 mg/L bromophenol blue) directly and proteins were isolated with 10% SDS-PAGE. The membrane was probed with anti-6×His mAb (Abcam) and anti-GAPDH pAb (Sangon), followed by incubation with HRP-conjugated anti-mouse IgG and anti-rabbit IgG (Santa Cruz Biotechology), respectively. Proteins were visualized with ECL.

### Luciferase activity assay

Cell lysate was prepared by using passive lysis buffer (Promega). Firefly and renilla luciferase activities (Fluc and Rluc) were measured using a Dual Luciferase Assay kit (Promega) according to the manufacturer’s instructions.

### Statistical analysis

All the experiments involving counting or calculation were performed independently at least three times and data are means ± standard deviation (SD). A Student’s two-tailed t test was used for statistical analysis by using GraphPad Prism 6.1 (GraphPad 6 Software, San Diego, CA). P< 0.05 was considered statistically significant.

## 3. Results

### 3.1 Expression of L3HYPDH inhibits EV71-GFP replication

To study the activity of L3HYPDH, the recipient cell line is best to have little or no IFN signaling in response to EV71 infection. We thus used HEK293A cell line as it lacks the expression of many pattern recognition receptors (PRRs) [23]. To support the infection, the predominant receptor SCARB2 was stably introduced into the cells (293A-SCARB2) [24]. To check the transfection efficiency, we co-transfected pCAG-DsRed with L3HYPDH-expressing plasmids in a mass ratio of 1:3. Under this biased ratio, we assumed that cells bearing fluorescent signals would also have L3HYPDH expressed. The transfected cells were then infected with EV71-eGFP 30 h later. Viral replication in the DsRed-positive populations was quantified by FACS assay of GFP at 18 h post infection (Figure 1A). Data showed that the GFP-positive population from empty vector-transfected cells was much larger than that from L3HYPDH-expressing cells (Figure 1B and 1C), suggesting L3HYPDH acts as an inhibitory host factor against EV71.

**Figure 1.**
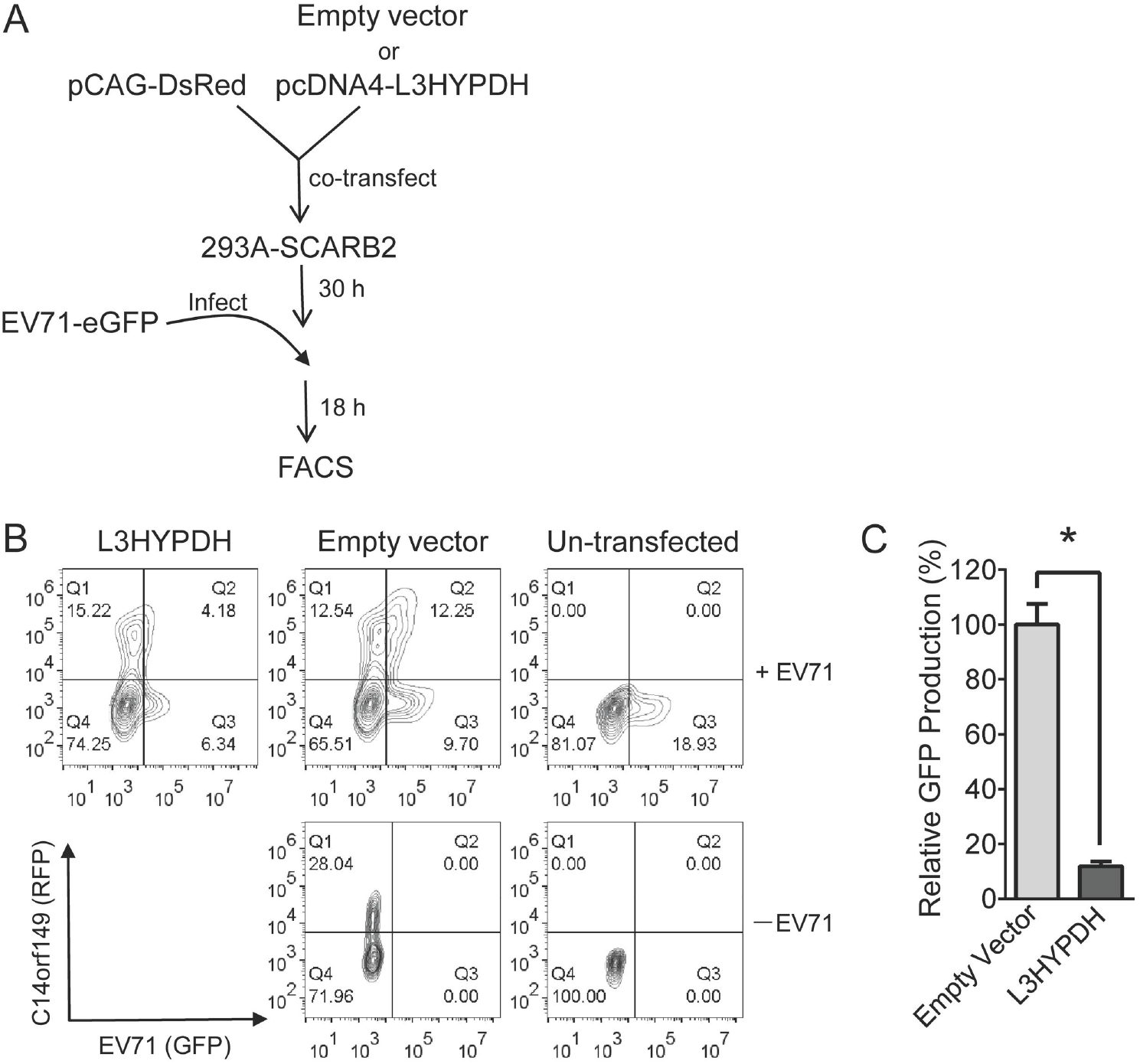
FACS-based assay for antiviral activity of L3HYPDH against EV71-GFP replication. (A) Overview of the procedures detecting anti-EV71 activity of over-expressed L3HYPDH using FACS. (B) FACS plots of L3HYPDH inhibition of EV71-GFP in 293A-SCARB2 cells. Numerals represent percent of total cell counts. (C) GFP production, which was calculated by multiplying GFP and DsRed co-positive cell number by the mean value of GPF intensity. The value of the control cells transfected with empty vector was set as 100%. Results are represented as means ± SD of three independent experiments. *, P< 0.05.

### 3.2 Expression of endogenous L3HYPDH suppresses EV71 replication

Next, we asked whether the endogenous L3HYPDH also inhibits EV71 infection. We set to test its expression pattern in commonly used cell lines. The RNA levels were measured by RT-qPCR. Data showed that HeLa cells express the least amount of L3HPDH while other tested cells have similar expression levels (Figure 2A). As is mentioned previously, this gene was identified as an ISG in immune cell lines, we then set to test if its expression is also responsive to IFN treatment here. Upon exposure to IFN-α2b, the level of L3HYPDH mRNA was up-regulated and peaked at about 18 h in 293A, 293A-SCARB2 and A549 and at about 12 h in Vero cells (Figure 2B). In contrast, the mRNA level changed little in HeLa cells and even decreased a little in RD cells (Figure 2B). The data also showed that SCARB2 expression in 293A cells (293A-SCARB2) does not change L3HYPDH expression either in unstimulated state or IFN-stimulated state.

**Figure 2.**
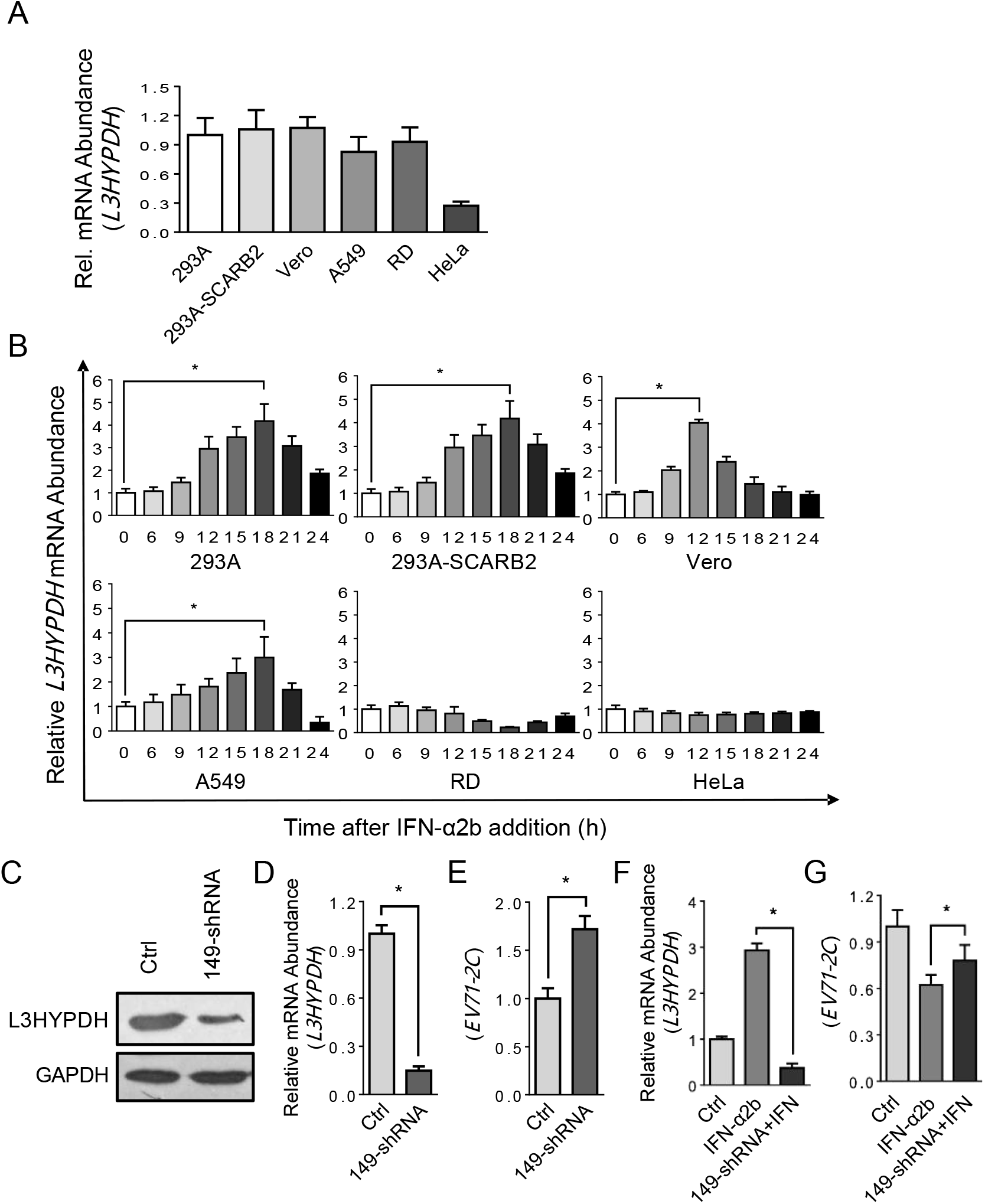
Anti-EV71 activity of the endogenous L3HYPDH. (A) RT-qPCR assay of the endogenous L3HYPDH mRNA level in different cell lines, which was normalized to GAPDH mRNA level. (B) RT-qPCR assay of L3HYPDH expression with IFN-α2b (1000 IU/ml) treatment for indicated time in different cell lines. L3HYPDH mRNA level was normalized to that of GAPDH. The relative mRNA level from untreated cells (marked as 0 h) was set as 1. (C) Western blot of knockdown efficiency of shRNA149 targeting L3HYPDH. 293A-SCARB2 cells were transfected with pcDNA4-L3HYPDH together with shRNA149-expressing plasmid or control plasmid (Ctrl) at a ratio of 1:3. The GFP-positive cells were sorted and then infected with EV71, followed by RT-qPCR analyses of L3HYPDH mRNA (D) and EV71 2C mRNA (E) after depression of L3HYPDH expression. RT-qPCR analyses of the L3HYPDH (F) and EV71-2C (G) mRNA in 293A-SCARB2 upon depression of L3HYPDH expression by shRNA in the presence of IFN-α2b. 293A-SCARB2 cells were seeded into a 10 cm dish and then transfected with 5 μg of pSUPER-GFP-shRNA149 or pSUPER-GFP. After incubation for 24 h, the GFP-positive cells were isolated using FACS and divided into two parts, one treated with 1000 IU/ml of IFN-α2b, the other mock-treated with water as control, followed by EV71-MZ infection (MOI, 0.1). Eighteen hours post-infection, total RNAs were isolated and used for RT-qPCR measurement. The target mRNA level was normalized to that of GAPDH, and the relative value from the control cells was set as 1. The results are represented as mean ±SD of three independent experiments. *, P<0.05.

We then tested the function of L3HYPDH using shRNA-mediated gene knockdown. An shRNA specific to L3HYPDH, designated as 149-shRNA, was designed and transcribed from pSuper-GFP-shRNA149. Its knocking down efficiency was detected in 293A-SCARB2 cells by co-transfecting pcDNA4-L3HYPDH together with pSUPER-GFP-shRNA149 or with pSUPER-GFP as a control. Western blot analysis showed that the myc-tagged L3HYPDH protein level decreased dramatically in the presence of shRNA149 (Figure 2C), suggesting this shRNA is potent in down-regulating the expression level of L3HYPDH. To determine if the endogenous L3HYPDH could suppress EV71 replication, 293A-SCARB2 cells were transfected with pSUPER-GFP-shRNA149, and the GFP-positive cells were sorted by FACS, followed by a clinical isolate of EV71, EV71-MZ infection. RT-qPCR assay revealed that 149-shRNA reduced L3HYPDH mRNA level by more than 80% (Figure 2D) and increased EV71-transcribed 2C mRNA level from 1 to 1.7 (Figure 2E), indicating that the expression of endogenous L3HYPDH impaired EV71 replication. Importantly, the L3HYPDH knockdown (Figure 2F) relieved IFN-mediated EV71 replication impairment (Figure 2G), suggesting both the basal level and IFN-stimulated expression of L3HYPDH encode antiviral activity against EV71.

### 3.3 Determination of the amino acid sequences essential for anti-EV71 activity of L3HYPDH

Next, we asked which domain is important for L3HYPDH antiviral activity. We mapped its regions using the FACS-based functional assay. Three N-terminal and three C-terminal serial deletions of L3HYPDH are schematically shown in Fig 3 (middle panel), designated as ΔN1, ΔN2, ΔN3, ΔC1, ΔC2, and ΔC3. Their coding sequences were cloned in fusion with a myc-6×His tag at the C-terminus as with the wild type (WT) L3HYPDH. The resulting plasmids were individually transfected into 293A-SCARB2 cells together with pCAG-DsRed followed by EV71-eGFP infection as described in Fig 1A. L3HYPDH protein expression was validated by western blotting (Figure 3, lower panel) and virus infection was detected by eGFP measurement (Figure 3, upper panel). FACS assay showed that L3HYPDH ΔN2 lacking the amino acids from position 1 to 120 significantly impaired the antiviral activity in comparison with WT, while L3HYPDH ΔN1 lacking the amino acids from position 1 to 60 only slightly weakened the antiviral activity. L3HYPDH ΔC1 lacking the C-terminal 60 amino acids from 295 to 354 also weaken the antiviral activity, while further deletion did not change this impairment. These results indicate that the amino acid sequences from position 61 to 120 and those from position 295 to 354 are both required for optimal anti-EV71 activity, and the N-terminus plays a more important role.

**Figure 3.**
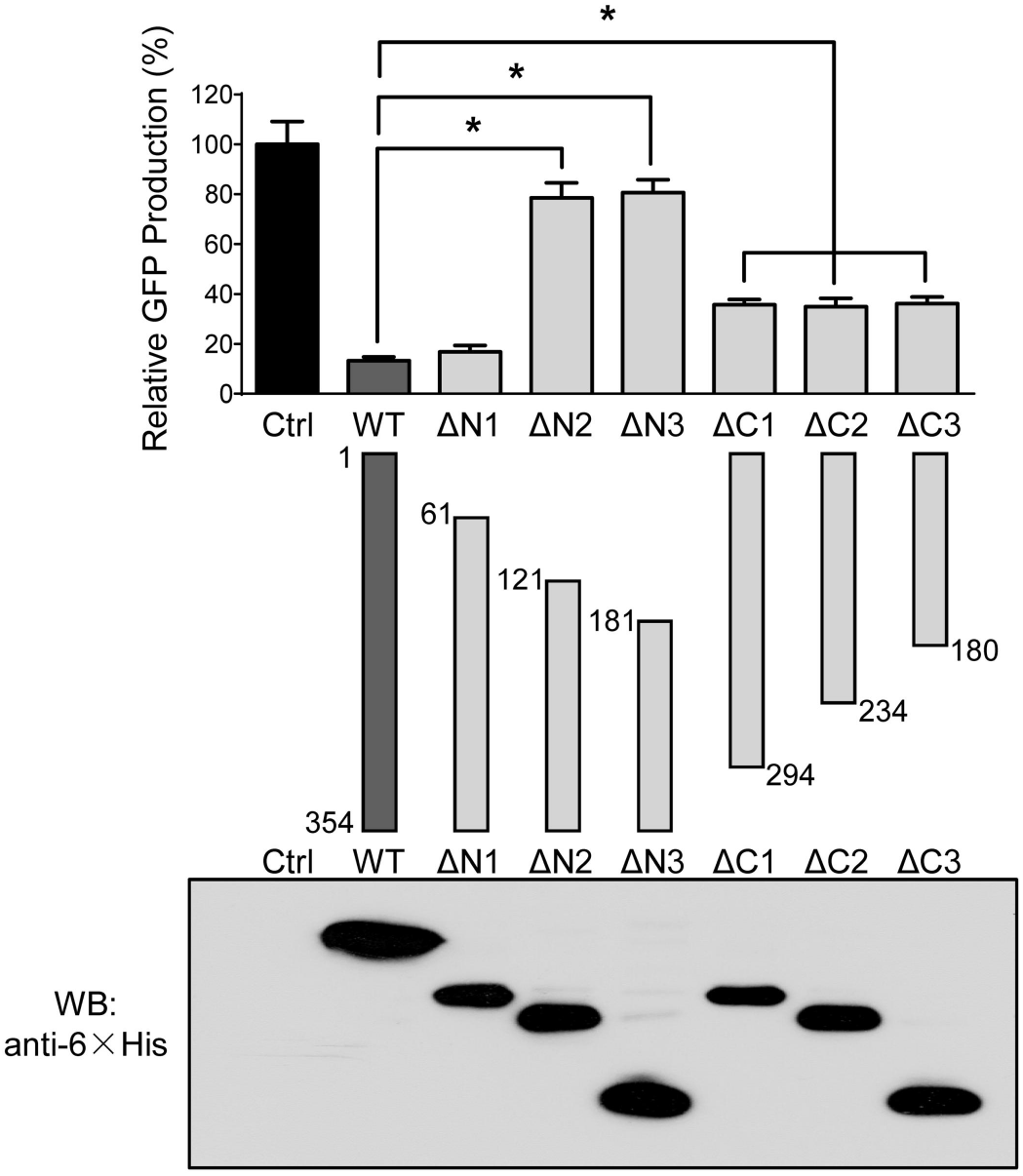
Mapping amino acid sequence required for anti-EV71 activity of L3HYPDH. Deletion mutants of L3HYPDH were schematically shown in the middle panel. Numbers indicate starting and ending amino acid. The plasmids expressing L3HYPDH wild type (WT) or truncated mutants were individually transfected into 293A-SCARB2 cells together with pCAG-DsREd at a ratio of 3:1, followed by EV71-GFP infection. Same performance was done the empty vector used as a control (Ctrl). L3HYPDH WT proteins or deletion mutants were analyzed by Western blot with anti-6×His mAb (lower panel). GFP production from EV71-GFP was detected using FACS and calculated as described in Fig1C. The value from control was set as 100%. Data are represented as mean ± SD of three independent experiments. *, P< 0.05.

### 3.4 EV71 replication is suppressed in the cell line 293A-SCARB2-L3HYPDH expressing L3HYPDH constitutively

A cell line stably expressing L3HYPDH (293A-SCARB2-L3HYPDH) was generated to facilitate the study of its mechanism of action. They were infected with EV71-eGFP at an MOI of 0.1. FACS assay showed that the eGFP production in 293A-SCARB2-L3HYPDH decreased significantly (Figure 4A). Upon infection with EV71-MZ, there was significantly less viral multiplication in 293A-SCARB2-L3HYPDH than in the control cells (Figure 4B). IFA showed that L3HYPDH proteins were mainly located in the cytoplasm (Figure 4C), consistent with its anti-EV71 action. Therefore, the cell 293A-SCARB2-L3HYPDH displays remarkable anti-EV71 activity due to the ectopic expression of L3HYPDH, and thus can be exploited to uncover the underlying antiviral mechanism.

**Figure 4.**
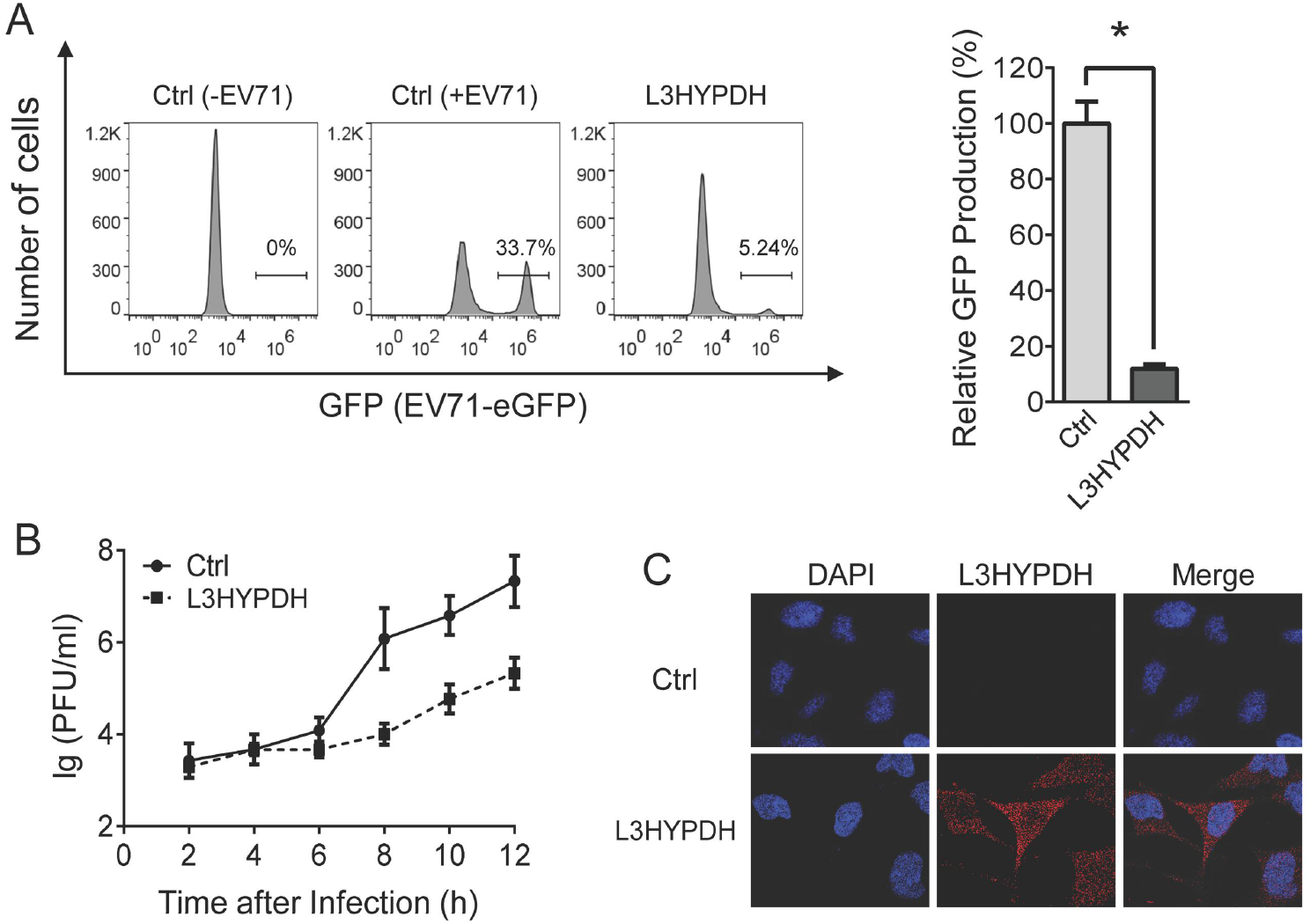
Evaluating antiviral activity of 293A-SCARB2-L3HYPDH cells. 293A-SCARB2-L3HYPDH and the control cell 293A-SCARB2-Ctrl were infected with EV71-GFP (MOI, 0.1). GFP production was detected using FACS (A) and calculated (B). Data are represented as mean ± SD of three independent experiments. *, P< 0.05. (C) Time-viral yield assay. 293A-SCARB2-L3HYPDH and the control cell were individually infected with EV71-MZ (MOI, 2). The culture supernatants were harvested at different time points as indicated and titrated by plague assay. Data for each time point are means ± SD of three independent experiments. (D) Subcellular localization of tagged L3HYPDH proteins using IFA. Nuclear DNA was stained with DAPI.

### 3.5 L3HYPDH interferes with the synthesis of viral RNA and proteins

The effects of L3HYPDH on different life stages of EV71 replication were examined in 293A-SCARB2-L3HYPDH cells. Based on the knowledge that EV71 is only adsorbed on the cell surface and could not finish endocytosis at 4°C, 293A-SCARB2-L3HYPDH and the control cells were incubated with EV71-MZ for 1 h at 4°C followed by IFA with the antibody specific to EV71 VP2. As shown in Fig 5A, massive number of virions distributed on the outer surfaces of both cell lines, showing no difference in numbers, indicating that L3HYPDH does not interfere with EV71 attachment. After attachment at 4°C, the viruses were further incubated with the cells for an additional 30 min at 37°C to complete endocytosis. IFA showed that viruses entered both cell lines with little difference (Figure 5B), indicating that L3HYPDH has no effect on EV71 endocytosis. In this way, these results demonstrate that L3HYPDH does not impede the viral attachment and endocytosis.

**Figure 5.**
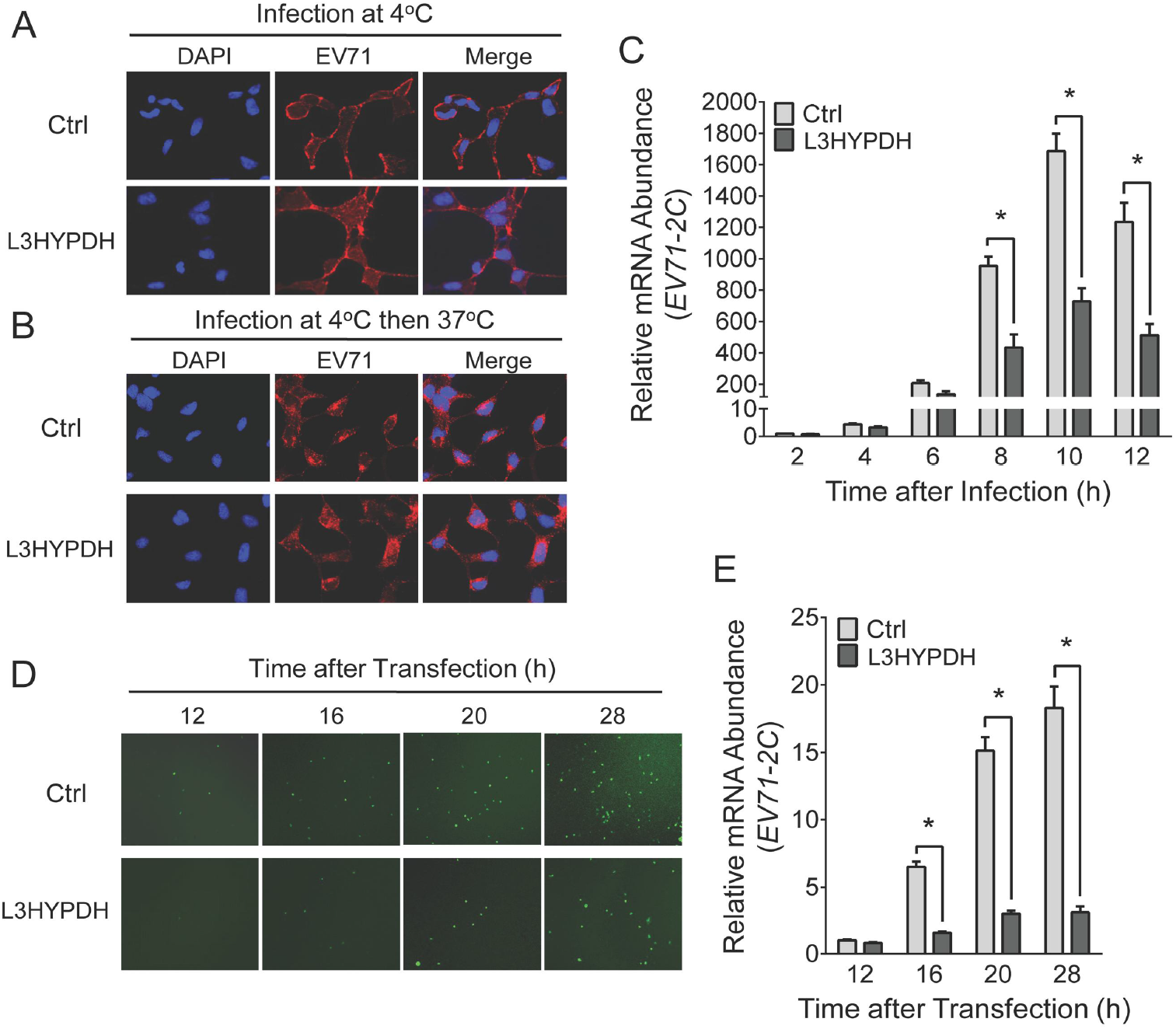
Stage assays for unveiling the mechanism of L3HYPDH against EV71. Effect of L3HYPDH on attachment (A) and endocytosis (B) of EV71 was examined using IFA. 293A-SCRB2-L3HYPDH and 293A-SCARB2-Ctrl cells were infected with EV71-MZ (MOI, 100). Nuclei were stained with DAPI. (C) Effect of L3HYPDH on viral RNA measured with RT-qPCR. 293A-SCARB2-L3HYPDH and control cell (Ctrl) were infected with EV71-MZ (MOI, 2). EV71 2C mRNA level was measured at indicated times and normalized to that of GAPDH, with the relative level in control cell at 2 h post infection set as 1. EV71-GFP RNAs were transfected into 293A-SCARB2-L3HYPDH and control cell (Ctrl). GFP signal and EV71 2C RNA level at different times post transfection were examined by fluorescent microscope (D) and RT-qPCR (E), respectively. All the RT-qPCR data are represented as means ± SD of three independent experiments. *, P<0.05.

The effects of L3HYPDH on the synthesis of viral RNA and proteins were investigated by monitoring changes in their levels over time. 293A-SCARB2-L3HYPDH and 293A-SCARB2-Ctrl cells were infected with EV71-MZ. Total RNAs were isolated at different times post-infection and the viral RNA abundance was measured by RT-qPCR. The increase in EV71 RNA levels over time in 293A-SCARB2-L3HYPDH cells was much lower than that in the control cells (Figure 5C). Due to the tight crosstalk between viral translation and viral RNA synthesis, we hypothesized that L3HYPDH might inhibit the synthesis of viral RNA, proteins, or both. To further confirm this assumption, EV71-GFP RNAs were transfected into 293A-SCARB2-L3HYPDH and the control cell, and then the viral RNA and GFP proteins were measured and compared at different times after transfection. Microscope and RT-qPCR analyses showed that both the number of GFP-positive cells and the viral RNA level were much lower in 293A-SCARB2-L3HYPDH cells than in the control cells (Figure 5D, 5E). These results suggested that L3HYPDH suppresses either EV71 RNA replication, viral protein synthesis, or both.

### 3.6 L3HYPDH impairs the translation mediated by EV71-5’UTR

The repression of L3HYPDH on viral protein synthesis was investigated using a bicistronic reporter system. As shown in Fig 6A, psiCHECK-2-based reporter plasmids were constructed with the HSV-TK promoter deleted to generate the control (psiCHECK2-M) or replaced with EV71-5’UTR or HCV-5’UTR, which contains EV71 IRES or HCV IRES, respectively. pcDNA4-L3HYPDH or the empty vector was transfected into 293A cells together with one of the three reporter plasmids at a ratio of 3:1, and then the luciferase activity and mRNA level were measured after incubation for 48 h. For these reporters, the mRNA level ratio of Fluc/Rluc in L3HYPDH-overexpressed cells was equal to that in the empty vector-transfected cells as revealed by RT-qPCR (Figure 6B). However, the luciferase activity ratio (Fluc/Rluc) showed variability (Figure 6C). Whether L3HYPDH was over-expressed or not, the Fluc/Rluc ratio of the control reporter was extremely low due to the absence of IRES; the ratio of the EV71-5’UTR-containing reporter reduced by 29% upon overexpression of L3HYPDH; while the ratio of the HCV-5’UTR-containing reporter changed little. We here proposed that L3HYPDH could specifically inhibit the reporter translation mediated by EV71 IRES. The knockdown assay further provided evidence for this speculation. 293A-SCARB2-L3HYPDH cells were transfected with the GFP-encoded 149-shRNA-expressing plasmid or the empty vector. Then the GFP-positive cells were isolated and transfected with the reporter plasmids. Compared to the control cells, the Fluc/Rluc ratio of the EV71-5’UTR-containing reporter in 293A-SCARB2-L3HYPDH cells increased upon L3HYPDH knockdown (Figure 6D). Altogether, these results indicate that L3HYPDH can specifically impair the translation initiated by EV71-5’UTR.

**Figure 6.**
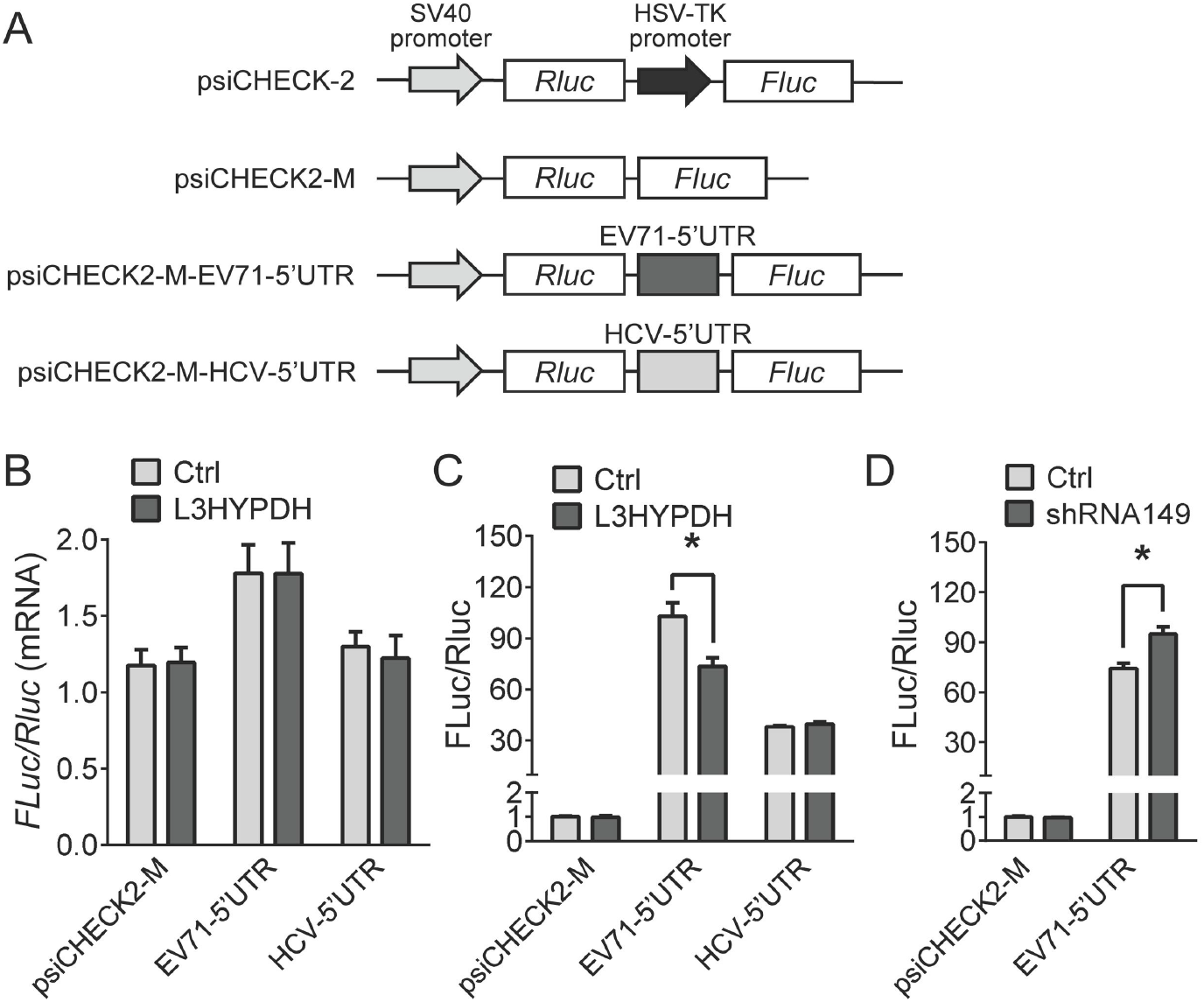
Bicistronic reporter assay to measure the effect of L3HYPDH on EV71 IRES mediated translation. (A) Schematic of bicistronic reporters. Rluc is translated in a cap-dependent manner and Fluc in an IRES-dependent manner. pcDNA4-L3HYPDH or empty vector was transfected into 293A cells together with psiCHECK2-M, psiCHECK2-EV71-5’UTR, or psiCHECK2-HCV-5’UTR. Effects of L3HYPDH on the reporter expression were estimated by RT-qPCR (B) and luciferase activity assay (C). Fluc/Rluc ratio was calculated, with the relative value from the cells transfected with empty vectors was set as 1. (D) RNAi assay of effect of L3HYPDH on reporter expression mediated by IRES. pSUPER-GFP-shRNA149 or pSUPER-GFP was transfected into 293A-SCARB2-L3HYPDH cell. The GFP positive cells were transfected with psiCHECK2-M or psiCHECK2-M-EV71-5’UTR. Luciferase activity was measured, with Fluc/Rluc ratio from cells co-transfected with psiCHECK2-M and pSUPER-GFP set as 1. Transfection with pcDNA4 was used as a control (Ctrl). Luciferase activities were measured. Fluc/Rluc ratio was calculated, with the relative value from the cells transfected with the empty vector set as 1. Data are means ± SD of three independent experiments. *, P< 0.05.

## 4. Discussion

In this work, we report that the ISG product L3HYPDH has antiviral activity against EV71 according to knockdown and over-expression experiments. Over-expression of L3HYPDH repressed eGFP production of EV71-eGFP (Figurer 1B) and caused significant inhibition of propagation of the clinical isolate EV71-MZ (Figurer 4B). Importantly, endogenous L3HYPDH is active in down-regulating EV71 mRNA in 293A-SCARB2 cells (Figurer 2E). Additionally, our data showed that IFN-α2b treatment was less effective against EV71 in cell culture when expression of L3HYPDH was depressed by shRNA (Figurer 2F, 2G). Therefore, L3HYPDH contributes to the antiviral activity of IFN-α2b against EV71. The potential activity of L3HYPDH against other viruses is not known. Given that different viruses are usually targeted by unique sets of ISGs [25], an extensive investigation on L3HYPDH will help to further elucidate the mechanism of IFN-mediated innate immunity against invading viruses.

Our data show that L3HYPDH may interfere with EV71 replication at post-entry stage (Figure 4). Bicistronic reporter assays confirmed that expression of L3HYPDH inhibited translation initiated by EV71 IRES (Figure 6B), however, the reporter protein was less reduced than EV71 RNA and virus-carrying eGFP production during the first round of infection (Figurer 1B, 4A, 5C-E). These inconsistences suggest that L3HYPDH hampers EV71 replication at other steps as well. Although inhibition of viral RNA replication is likely, other potential effects on viral RNA stability, viral assembly and viral release cannot be excluded. Therefore, L3HYPDH inhibits EV71 replication at least at two levels, and these data agree with previous studies indicating that many ISGs block viral replication at multiple stages of the viral life cycle [26]. Considering that a range of proteins are involved in the viral RNA replication and translation process, we performed co-immunoprecipitation and tandem affinity purification combination mass spectrometry to screen for proteins interacting with L3HYPDH. Neither viral nor host proteins were identified (data not shown). These results suggest that the association of L3HYPDH proteins with other proteins should be transient or weak. L3HYPDH might also function by binding to the viral RNA directly; however, no known RNA-binding domains were predicted with online software (data not shown).

Viral translation is in most cases host cell dependent. To maximize efficiency, different viruses evolved many strategies to facilitate selective translation of viral mRNAs over host transcripts [3,27]. Among these, the IRES-mediated translation initiation is necessary for picornavirus and hepacivirus to replicate [4,28]. Reporter assays showed that expression of L3HYPDH impaired initiation of translation mediated by EV71 IRES but not HCV IRES (Figurer 6B, 6C). These two tested IRES differ in nucleotide length and structure as well as in host factors required for translation initiation and regulation [29]. A potential target of L3HYPDH should be involved in EV71-5’UTR-mediated translation. Meanwhile, despite being present in all picornaviruses, IRES is also diverse in length and structure and requires different host factors to function [30,31]. Whether L3HYPDH can inhibit other genuses of picornavirus by interfering with IRES-mediated translation is not clear is not clear, but this is worthy of study in the future.

L3HYPDH is a trans-3-hydroxy-L-proline dehydratase, and specifically catalyzes the dehydration of dietary trans-3-hydroxy-L-proline and from degradation of proteins such as collagen IV that contain it. This dehydratase contains two active sites, a Cys residue at the 104 position and a Thr residue at the 273 position [15]. Interestingly, the region required for anti-EV71 activity was mainly mapped to the amino acid sequence from position 61 to 120 of L3HYPDH protein (Figure 3), which contains the Cys104 active site. Whether this proline dehydratase activity is involved in the anti-EV71 activity is not known yet, but at least an extra region from the C-terminus is required for L3HYPDH antiviral activity.

In sum, our current work uncovered a new anti-EV71 host factor whose expression is up-regulated by IFN-I treatment in certain cell lines, highlighting the protective role of IFN-I and ISGs upon viral infection. Understanding ISG products and antiviral spectra, as well as their mechanisms of action and biological function will help create novel therapeutics for HFMD in the future.

## Author Contributions

X.G, J.L, L.L, S.Z, X.Meng, N.L, H.Y, R.L, and X.Mu designed the assays. J.L, L.L, S.Z, X.Meng, N.L, H.Y, R.L did experiments. X.G, J.L, L.L, and X.Mu analyzed the data and wrote the manuscript.

## Data Availability Statement

Source data can be obtained upon reasonable request to the corresponding authors.

## Acknowledgments

We thank Dr. Liguo Zhang for providing the plasmids pCAG-DsRed, pWSK-EV71-GFP, pcDNA3.1-T7RNP and the cell line 293A-SCARB2. We also thank Dr. Guangxia Gao for providing the plasmids pLPCX-C14orf149 and pNL4-3RL-HCV-FL.

