## Supplemental Table 1 for "Inhibition of EV71 replication by an interferon-stimulated gene product L3HYPDH"

Table S1 Primers used in this study

| **Name** | **Sequence (5’-3’)** | **Target gene, usage** |
| --- | --- | --- |
| L3HYPDH-F | CGGGATCCGCCACCATGGAGAGCGC^a^ | *L3HYPDH* |
| L3HYPDH-R | CACGCGGCCGCACTTGAGAAGAAATCCA |  |
| L3HYPDH-N1-F | CACGGTACCATGGTGCGGCGACGGCTCA | L3HYPDHΔN1 |
| L3HYPDH-N-R | CACTCTAGACTGCACTTGAGAAGAAAT |  |
| L3HYPDH-N2-F | CACGGTACCATGGTGCCGGCGCCCCCTG | Pairing with L3HYPDH-N-R, L3HYPDHΔN2 |
| L3HYPDH-N3-F | CACGGTACCATGGGAAAGGTGATGGTGG | Pairing with L3HYPDH-N-R, L3HYPDHΔN3 |
| L3HYPDH-C-F | CACGGTACCATGGAGAGCGCGCTGG | L3HYPDHΔC1 |
| L3HYPDH-C1-R | CACTCTAGACTCATCTGGTTCAGTTCC |  |
| L3HYPDH-C2-R | CACTCTAGACTTTCACTATCAGGATGA | Pairing with L3HYPDH-C-F, L3HYPDHΔC2 |
| L3HYPDH-C3-R | CACTCTAGACTATGTCCAGGAACATCC | Pairing with L3HYPDH-C-F, L3HYPDHΔC3 |
| CHECK2-F | CACGCGGCCGCTCTAGGTTTAAA | Back-to-back primers, used for generating psiCHECK2-M |
| CHECK2-R | CACGTCGACATGGCCGATGCTAAGAACATTA |  |
| EV71-5’UTR-F | CACGCGGCCGCTTAAAACAGCCTGTGG | EV71-5’UTR |
| EV71-5’UTR-R | CACGTCGACGTTTAGCTGTGTTAAGG |  |
| HCV-5’UTR-F | CACGCGGCCGCGGCGACACTCCACCATAG | HCV-5’UTR |
| HCV-5’UTR-R | CACGTCGACGATGCACGGTCTACGA |  |
| QL3HYPDH-F | AGGAGTGACAGCC CGAATTG | *L3HYPDH*, used for qPCR |
| QL3HYPDH-R | CACATTTCGCTTCCCTCACAG |  |
| QEV71-2C-F | TGTATGTCTCATTATCAGGGG | EV71 *2C*, used for qPCR |
| QEV71-2C-R | CCACCTGTTGCTTGTAACCGT |  |
| QRluc-F | ATAACTGGTCCGCAGTGGTG | *Rluc*, used for qPCR |
| QRluc-R | AGGCC GCGTTACCATGTAAA |  |
| QFluc-F | AGCACTTCTTCATCGTGGACCG | *Fluc*, used for qPCR |
| QFluc-R | GGCAGCTCGCCGGCATCGTCGT |  |
| H-QGAPDH-F | GAAGGTGAAGGTCGGAGT | Human *GAPDH*, used for qPCR^b^ |
| H-QGAPDH-R | GAAGATGGTGATGGGATTTC |  |
| Vero-QGAPDH-R | GAAGATGGTGATGGGGCTTC | Pairing with H-QGAPDH-F, Monkey *GAPDH*, used for qPCR of the RNA from Vero cells |

^a^ Restriction sites are underlined.

^b^The primers against human *GAPDH* have been described elsewhere . All other primers used for the PCR assays are designed using the Primer-BLAST tool (https://www.ncbi.nlm.nih.gov/tools/primer-blast/) or acquired from Primer Bank (https://pga.mgh.harvard.edu/primerbank/).
